# Cell and tissue-specific glycosylation pathways informed by single-cell transcriptomics

**DOI:** 10.1101/2023.09.26.559616

**Authors:** Panagiotis Chrysinas, Shriramprasad Venkatesan, Isaac Ang, Vishnu Ghosh, Changyou Chen, Sriram Neelamegham, Rudiyanto Gunawan

## Abstract

While single cell studies have made significant impacts in various subfields of biology, they lag in the Glycosciences. To address this gap, we analyzed single-cell glycogene expressions in the Tabula Sapiens dataset of human tissues and cell types using a recent glycosylation-specific gene ontology (GlycoEnzOnto). At the median sequencing (count) depth, ∼40-50 out of 400 glycogenes were detected in individual cells. Upon increasing the sequencing depth, the number of detectable glycogenes saturates at ∼200 glycogenes, suggesting that the average human cell expresses about half of the glycogene repertoire. Hierarchies in glycogene and glycopathway expressions emerged from our analysis: nucleotide-sugar synthesis and transport exhibited the highest gene expressions, followed by genes for core enzymes, glycan modification and extensions, and finally terminal modifications. Interestingly, the same cell types showed variable glycopathway expressions based on their organ or tissue origin, suggesting nuanced cell- and tissue-specific glycosylation patterns. Probing deeper into the transcription factors (TFs) of glycogenes, we identified distinct groupings of TFs controlling different aspects of glycosylation: core biosynthesis, terminal modifications, etc. We present webtools to explore the interconnections across glycogenes, glycopathways, and TFs regulating glycosylation in human cell/tissue types. Overall, the study presents an overview of glycosylation across multiple human organ systems.

## INTRODUCTION

Glycosylation is a ubiquitous post-translational modification that results in the formation of an array of cellular complex carbohydrate structures or glycans (1). These glycans, which appear either in branched or extended form on the cell surface or as single-monosaccharide additions within cells, control or fine-tune a multitude of biological functions during normal physiology and disease (2,3). The common glycoconjugate types on mammalian cells include the branched N-linked glycans on glycoproteins, O-GalNAc (N-Acetyl Galactosamine) type O-glycan modifications on glycoproteins, long repeating saccharide chains called glycosaminoglycans (GAGs) on a select set of proteoglycans, carbohydrate modifications on glycolipids, and finally O-GlcNAc (N-Acetyl Glucosamine) type single residue modifications on nuclear proteins and transcription factors. Besides these major families of glycoconjugates, glycans also form the anchor for Glycosylphosphatidylinositol (GPI)-linked cell-surface proteins. There also exist a growing list of rarer O-linked glycan modifications, including O-Glc (Glucose), O-Fuc (Fucose) and O-Man (Mannose) type glycosylation (4).

Glycans on cells are formed by the concerted action of ∼2% of the expressed proteome that are collectively called ‘glycoEnzymes’. These enzymes are products of the corresponding ‘glycogenes’. An ontology called “GlycoEnzOnto” has recently been curated to describe the existing knowledge of human glycoEnzymes within the domain of Glycosciences (5). In this ontology, the ∼400 glycogenes are annotated according to their molecular functions, biological processes, and physical location (cellular component), following the Gene Ontology convention (6). Besides the genes encoding enzymes in the glycan biosynthesis, GlycoEnzOnto also includes the entities involved in the regulation of nucleotide-sugar metabolism, glycosyl-substrate/donor transport, glycan degradation and other regulatory components. From the molecular function perspective, glycogenes are grouped into glycosyltransferases, other transferases (e.g., sulfotransferases), modifying enzymes (e.g., epimerases and kinases), glycosidases, molecular transporters and other regulators. Additionally, glycogenes/glycoEnzymes can also be classified according to their role in glycoconjugate biosynthesis, including: (i) The ‘initiation’ step that results in the attachment of the first monosaccharide or oligosaccharide to the protein/lipid; (ii) The ‘elongation and branching’ reactions that extend the original glycan often via lactosamine chain synthesis/branching; and (iii) The ‘termination or capping’ processes that prevent further chain extension. The goal of GlycoEnzOnto was to provide a shared resource that can be used for Systems Glycobiology analysis (7–11).

This study harnesses single-cell transcriptome data of humans and the aforementioned GlycoEnzOnto to shed light on the variation of glycopathway expression across various cell and tissue types. In particular, it analyzes the Tabula Sapiens (TS) dataset that generated single-cell RNA-sequencing (scRNA-seq) data for 483,152 human cells from 15 donors which had been organized into 24 tissues and over 400 cell types (12). Such single-cell data identify subtle, yet potentially vital, differences in glycosylation processes among different cells within the same tissue or organ as this is not possible using bulk sequencing. Our results show that only ∼50% of glycogenes are expressed in a given cell, with expression levels varying depending on gene function. In contrast to conventional thinking based on microarray/quantitative-PCR data analysis (13) that suggests that glycogenes are lowly expressed, our more holistic single-cell analysis reveals that the glycogenes are expressed at levels that are comparable to other protein-coding genes. Further, our analysis presents a map of glycopathway expression across tissue, illustrating the inherent heterogeneity across human cells and tissues. Specifically, the findings showed how enzymes involved in the metabolism of nucleotide sugars, glycan degradation processes, and biosynthesis of core structures, exhibit uniform and ubiquitous expression patterns across cell types and tissues, consistent with their foundational roles in glycan biosynthesis. Meanwhile, terminal glycoenzymes often serve specialized roles, and they are more selectively expressed in individual cell types. Lastly, the analysis of transcriptional factors using mutual information of TF-glycogene expression in the TS dataset fills the gap in knowledge of transcriptional regulators of glycosylation (14). The result reveals five regulatory modules, with each module controlling a different aspect of glycosylation. To bolster accessibility, we also developed webtools that allow further exploration of glycogenes, glycopathways and related transcription factors at single cell level: http://vgdev.cedar.buffalo.edu/glycocarta/ and http://vgdev.cedar.buffalo.edu/glycotf/.

## MATERIAL AND METHODS

### Data preprocessing

The scRNA-seq data in the Tabula Sapiens (TS) project were generated using two different single-cell sequencing technologies: 10X and Smart-seq, with the majority of the data coming from 10X (456,101 cells vs. 27,051 cells). Tabula Sapiens scRNA-seq data were obtained from the public website (15). This study focused only on scRNA-seq data from 10X platform to avoid any potential batch effects associated with different sequencing platforms.

The data preprocessing is illustrated in **Fig. 1A**, with individual steps being carried out using the Python package scanpy (16). The analysis started with the decontaminated UMI counts from the TS dataset. Decontamination of background RNA was previously performed using the method decontX (17). Following the standard practice, UMI counts were scaled cellwise so that each cell has a count depth of 10,000. This scaling produced relative RNA abundances 𝑥_!_ that are comparable across cells.

**Figure 1.**
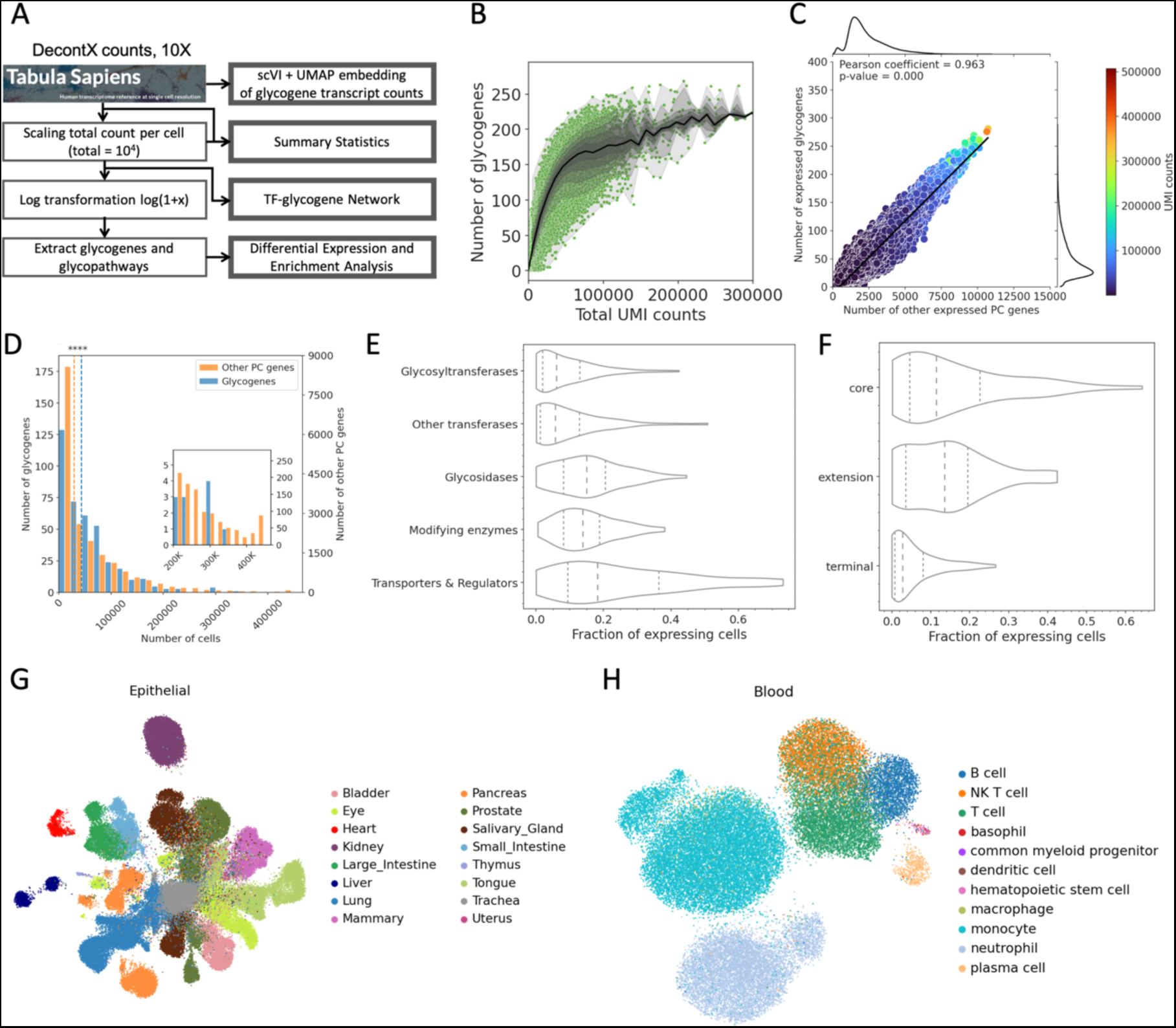
Transcriptomic analysis of glycogenes at single cell level. (A) Summary of data preprocessing and analyses of Tabula Sapiens (TS) data (see Methods for details). (B) Number of glycogenes with RNA count > 0. Each dot represents a cell. Dark line represents the median value. The shaded areas show contour of percentiles at 10 percent increments from the median, spanning 0^th^ to 100^th^ percentile. The data collection results in a shoulder at ∼65,000 UMI counts/cell corresponding to ∼165 glycogenes. (C) A positive correlation is observed between the number of detected glycogenes vs. other protein coding (PC) genes. The face color represents the UMI depth. (D) Distribution of PC and glycogene expressions in terms of the number of expressing cells (*i.e.*, cells with nonzero RNA count for the gene). Glycogenes are generally more commonly expressed in the TS cells than other PC genes. (E) Distribution of single cell expression of glycogenes. Glycogenes are grouped based on their biological functions as defined in the GlycoEnzOnto (see **Supplementary Table S2**). ‘Transporters and Regulators’ are generally more broadly expressed compared to other glycogenes including glycosyltransferases. (F) Distribution of single cell expression of glycosyltransferases in Core, Extension and Terminal groups (see **Supplementary Table S3**). Core enzyme expression is higher compared to extension and terminal modifiers (G-H) UMAP visualization of scVI latent embedding of glycogene expression in epithelial cells and blood tissue.

For differential expression analysis, the scaled UMI counts were log-transformed (*i.e.*, log (𝑥 + 1)) to satisfy the input requirement of the method MAST (18). For subsequent analyses, we subset the preprocessed count matrix to 19,847 transcripts (from a total of 58,559) associated with protein-coding (PC) genes as defined in BioMart. On average, PC genes make up 89.4% of the total number of reads. Out of these, we extracted data for 400 glycosylation-related genes (glycogenes) as defined in the GlycoEnzOnto (5). Python and R codes used for data analysis are available from: http://www.github.com/cabsel/glycots.

### scVI+UMAP embedding

To visualize glycogene expression in single cells, Single-cell Variational inference (scVI) was applied to the decontaminated glycogene UMI counts to generate a lower dimensional embedding of the glycogene expression (19). The method scVI produces a probabilistic latent space of single-cell gene expression data based on zero-inflated negative binomial distribution. Specifically, the Python’s scvi-tools package (20) was implemented with the following parameters: n_latent_ = 50, n_layers_ = 3, n_latent_ = 50, and dropout rate = 50. For this study, only the variational posterior of the scVI model was employed. For visualization of this embedding, a Uniform Manifold Approximation and Projection (UMAP) using n_neighbors_ = 15 and n_components_ = 2 (2D) was used (21).

### Differential expression analysis of glycopathways

Glycosylation-related pathways (glycopathways) are described in **Suppl. Table S3**. DE analysis using MAST requires as inputs the log(𝑥 + 1)-transformed UMI counts for the glycogene expression. For each pathway, its expression was evaluated by averaging the expression of glycogenes belonging to that pathway. Differential expression (DE) analysis of glycopathways was then performed using the MAST (Model-based Analysis of Single cell Transcriptomics). MAST adapts a hurdle model to tackle zero-inflation and biomodality of single-cell transcriptome data (18). As illustrated in **Fig. 2A**, MAST produces fold-change differences of the mean expression of a glycopathway between cells from a specific tissue, with respect to all other tissue with the same glycopathway. Associated statistical significance is also calculated. The DE analysis was implemented using the Seurat package (version 4.1.0) in R, specifically *FindAllMarkers* function (22).

**Figure 2.**
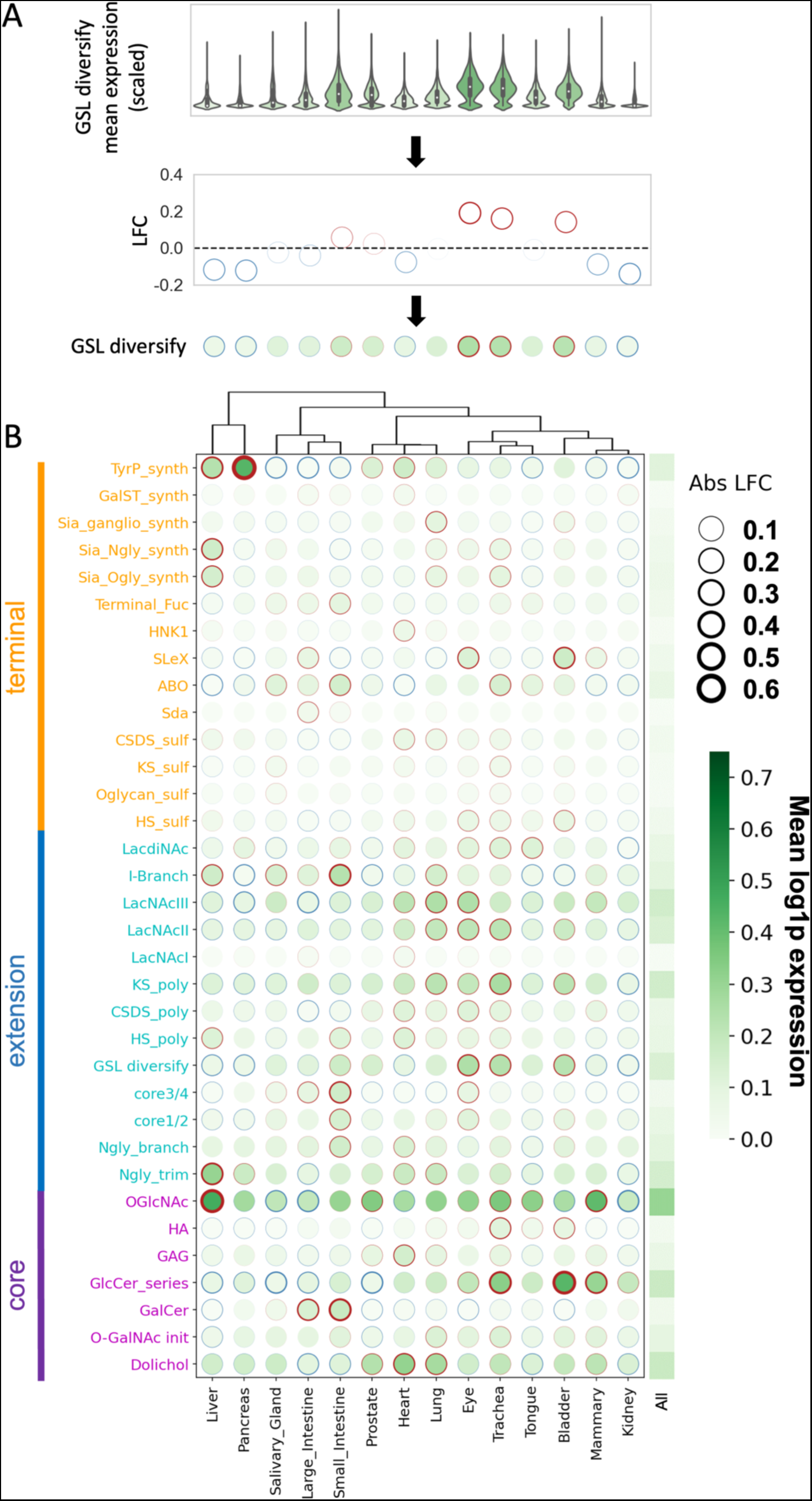
Differential expression (DE) analysis of glycopathway in epithelial cells. (A) “GSL (glycosphingolipids) diversify” pathway including the gene set B4GALNT1, B3GALT4, B3GNT5, B3GALT5, B4GALT1, B3GALNT1 and A4GALT, is used to illustrate the calculation scheme. Here, mean pathway gene expression in each tissue is first calculated from the zero-inflated single-cell data. Log-fold change (LogFC) expression is then determined, and this is presented using dots where a thicker red (blue) linewidth represents higher (lower) levels of the pathway expression in a given tissue with respect to all other tissue. The face color of the dot (greyscale) represents the mean expression of the glycopathway among the cells in a specific tissue. (B) DE of glycopathways is presented for selected Core, Extension and Terminal pathways for epithelial cells. The greyscale heatmap in the last column presents the mean glycopathway expression among all epithelial cells. As an example, O-GlcNAc related genes (OGA, OGT) are highly expressed across tissue. Among the tissue, this is most highly expressed in the liver compared to kidney and large intestine. In contrast the GSL diversity genes are higher in eye and trachea compared to other tissue.

### Glycopathway enrichment analysis

Enrichment analysis was performed to assess whether cells from a specific tissue are over-represented or depleted with cells expressing a given glycopathway. Here, a cell is labeled as an “expressing cell” when the average expression (scaled UMI count) of genes in a glycopathway is nonzero. For a given pair of glycopathway and tissue, a contingency table was constructed to distribute the cells into two distinct categorizations: expressing cells / non-expressing cells and cells in the tissue / cells not in the tissue, as follow:

|  | Expressing cells | Non-expressing cells |
| --- | --- | --- |
| Cells in tissue | $a$ | $b$ |
| Cells not in tissue | $c$ | $d$ |

The odds ratio OR, given by 𝑎𝑑⁄𝑏𝑐, indicates the overrepresentation (odds ratio > 1 or log(OR) > 0) or depletion (odds ratio < 1 or log(OR) < 0) of expressing cells in a tissue. The statistical significance was established via Fisher’s exact test based on hypergeometric sampling. The enrichment analysis was implemented in Python using *fisher_exact* function from the scipy package (version 1.10.1).

### Transcriptional factor analysis

The curation of TF-glycogene and glycopathway interactions involved evaluating mutual information (MI) of single-cell expression between every possible pair of TF-glycogene in the TFLink database (23). TF-gene interactions in the TFLink database were originally compiled from numerous databases that relied on different evidence of TF binding on the regulatory elements of the genes (23). Here, MI was used to provide additional evidence for TF-glycogene interactions based on shared information in their single-cell expression. The following equation gives the basis for evaluating MI for TF-glycogene relations:

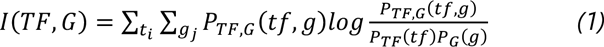

where 𝑃_𝑇𝐹,𝐺_ denotes the joint probability distribution of single-cell expression of a transcription factor 𝑇𝐹 and a glycogene 𝐺, and 𝑃_𝑇𝐹_ and 𝑃_𝐺_ denote the marginal probability distribution of 𝑇𝐹 and 𝐺, respectively. We evaluated 𝐼(𝑇𝐹, 𝐺) using the scaled UMI counts from all 10X cells in the TS dataset. The calculation of MI was performed using the Python package sklearn (24).

### Transcriptional Regulatory Module analysis

Transcriptional regulatory modules were identified by hierarchical clustering of glycogenes based on the log1p of MI, i.e. log(1+MI). Hierarchical clustering was performed using the method ‘complete-linkage’ to produce the distance matrix. Subsequently, the clusters of glycogenes (n = 5) were obtained using *fcluster* function in the scikit-learn package (version 1.2.2) with the criterion ‘maxclust’.

Each glycogene cluster was taken as a transcriptional regulatory module. Enrichment analysis was performed using the Fisher’s exact test to identify the over-representation of glycopathways. For this purpose, we employed the glycopathway definition given in **Suppl. Table S3**. Given two sets of glycogenes, one from a TRM and another from a glycopathway, we constructed the following contingency table:

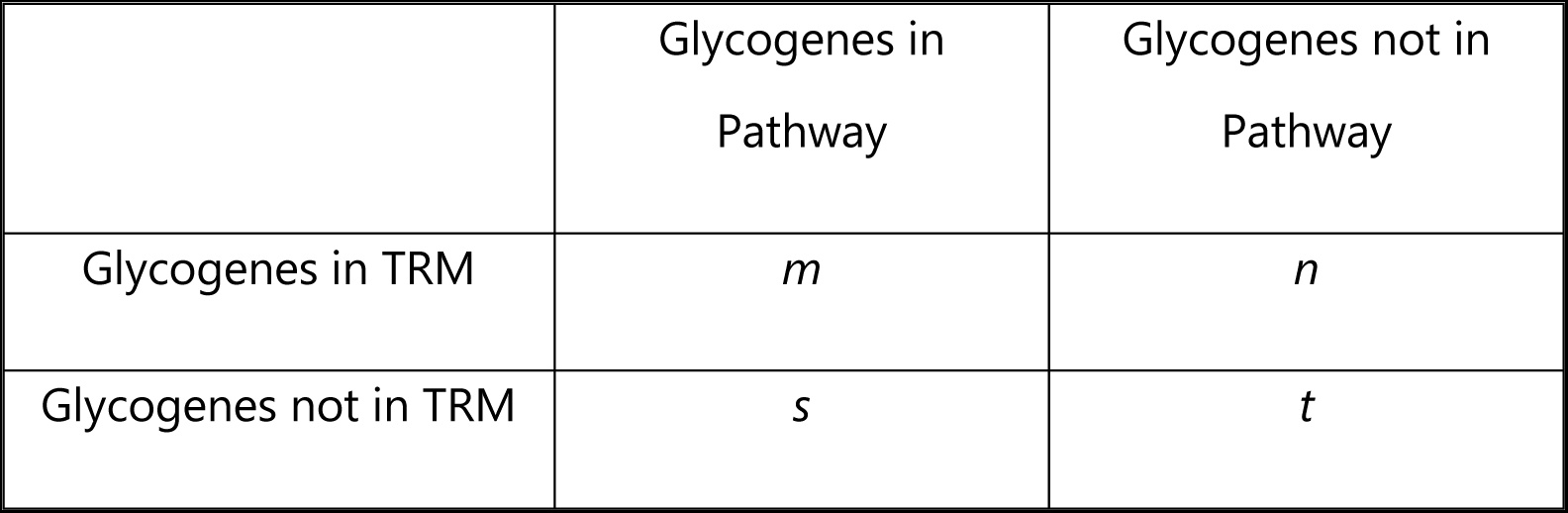

The odds ratio calculation (𝑂𝑅 = 𝑚𝑡⁄𝑛𝑠) and Fisher’s exact test were performed following the same procedure for the ED analysis (see **Glycopathway enrichment analysis**). Hierarchical and *k-*means clustering were implemented using the scikit-learn package (version 1.2.2) and the Fisher’s exact test using the scipy package (version 1.10.1) in Python. The statistical significance was established using Benjamini-Hochberg adjusted *p*-values to account for multiple hypothesis tests (25).

To identify the key TFs, we developed a ranking procedure that generates a list of TFs, ordered based on their relevance to each TRM. First, for every glycogene within a cluster, we sorted the TFs according to their MI score with the glycogene. Then, we calculated an average ranking for each TF across all glycogenes within the cluster. This process was replicated for each glycogene cluster. The results of this analysis are presented in **Suppl. Table S6.**

## RESULTS

### Overview of glycogene expression in single cell studies

Single-cell RNA-sequencing (scRNA-seq) technologies are transforming our understanding of cell biology, from development (26,27) to immune systems (28,29) and to aging (30,31). While different single-cell methods have their advantages and disadvantages, they share a few issues, such as low mRNA capture efficiency and high dropout rates, that particularly affect the measurement of genes with low expression (32,33). As glycogenes are traditionally thought to be lowly expressed (13), this study assessed the ability of single-cell data to inform us about genes and pathways involved in glycosylation by analyzing scRNA-seq data from the Tabula Sapiens (TS) project.

We applied the bioinformatics analysis workflow depicted in **Fig. 1A** to the TS dataset, focusing on glycogenes and glycopathways presented in GlycoEnzOnto (5). We compared the expression profiles of 400 glycogenes to 19,447 protein coding (PC) genes in the TS dataset (**Suppl. Table S1)**. As expected, the number of detectable glycogenes in the TS cells increased with the cell’s UMI count (**Fig. 1B**). Specifically, at the median (mean) UMI count depth of 6496 (10181) we detected between 2 and 98 glycogenes (2 to 118 genes) with RNA count > 0 and a median value of 40 glycogenes/cell (mean value of 58 glycogenes/cell). Thus, only 10% of all glycogenes are detected in the median cell in the TS dataset (14.5% of glycogenes for the mean cell). The low level of detection of glycogenes (*i.e.*, 10-14.5%) might stem from the overall low expression of glycogenes. In this regard, increasing UMI count depth did enhance glycogene detectability. However, the maximum number of captured glycogenes reached a plateau at ∼220, suggesting that, at most, only 50-60% of all glycogenes are expressed in individual cells. Lastly, only a marginal improvement in the number of detected glycogenes was observed beyond ∼65,000 UMI counts per cell, at which point about 165 glycogenes (median) were detected (**Fig. 1B**).

We observed a strong correlation between the number of glycogenes and the number for other PC genes detected in cells (**Fig. 1C**). Comparing the distribution of expression between glycogenes and other PC genes, using the fraction of expressing cells as an indicator of gene expression level, revealed a significant difference between the two groups (*p-*value = 1.19×10^−7^, Kolmogorov-Smirnov test) (see **Methods** and **Fig. 1D**). Interestingly, in the TS cells, glycogenes were more commonly expressed than other PC genes (p-value = 5.33×10^−5^, two-sided Wilcoxon ranksum test). But, despite this prevalence, glycogenes were not among the highly expressed genes in the TS dataset (**Fig. 1D** inset)—in fact, glycogenes were depleted among the top 10% of highest expressing PC genes (odds ratio = 0.655, p-value = 0.03, two-sided Fisher exact test). Overall, the data suggested that while the glycogenes may not be among the most highly expressed genes, they are ubiquitously expressed commonly at levels comparable to or higher than the average PC gene.

### Variability in glycogene expression patterns across cell types and tissues

We delved deeper into the variability of glycogene expression among functional sub-groups using the GlycoEnzOnto as a guide. To do this, we evaluated the fraction of cells expressing glycogenes across different sub-groups (**Suppl. Tables S2 and S3**). **Fig. 1E** reveals that glycogenes belonging to the ‘Transporters and Regulators’ sub-groups—that is, genes involved in the creation of nucleotide-sugars, monosaccharide transport and related metabolism—generally exhibited higher expression than other glycogene sub-groups. Glycogenes responsible for glycan modifications and those producing glycosidases displayed comparable expression levels with both gene groups presenting moderate expression. Finally, the glycotransferases and other transferases demonstrated the lowest expression levels. Delving further into glycogenes associated with the biosynthesis of core structures (Core), glycan chain elongation (Extension), and capping of glycan structures (Terminal), the Core and Extension groups showed similar levels of expression that were higher than the expression of glycogenes in the Terminal group (**Fig. 1F**). These patterns align with the role of the core enzymes in initiating the formation of specific glycan types, except perhaps for the case of O-GalNAc type carbohydrate chain formation that can be catalyzed by various isoenzymes. Thus, these Core genes are more broadly expressed in various cells compared to terminal modifiers that are expressed in a tissue specific manner (34). Although our analysis employed the fraction of expressing cells as the metric for gene expression level—following the recommendation for lowly expressed genes (35)—we observed similar trends using the mean expression of genes across cells (see **Suppl. Fig. S1**).

Next, we investigated how glycogene expression pattern in individual cells varies across different cell types and tissue types using scVI (single-cell Variational Inference) for latent embedding and UMAP (Unified Manifold Approximation and Projection) for 2D visualization (**Fig. 1G and H**, and **Suppl. Fig. S2**). Examining epithelial cells (**Fig. 1G**), clusters (grouping) of cells emerged in the UMAP plot following their tissue sources. Interestingly, even for cells from the same tissue, for example liver, pancreas and salivary gland, distinct cell groupings appeared. We made similar observations for endothelial, stromal and immune cells in the TS dataset (see **Supplementary Figure S2A-B**). Shifting focus to cells in the blood tissue (**Fig. 1H** and **Fig. S2D**), these cells formed clusters according to their lineage along the hematopoietic stem cell (HSC) differentiation pathway. Specifically, cells from the lymphoid path, including B-cells, T-cells, Natural Killer (NK) T cells, and plasma cells appeared in overlapping clusters, while cells from the granulocyte-macrophage lineage (macrophages, monocytes, and neutrophils) formed separate groups. The overt grouping of cells, influenced by their tissue of origin and lineage, suggests that mammalian tissues and cell types possess unique single-cell glycogene expression patterns, potentially indicating their varied glycan structures. In subsequent analyses, as 2D UMAP plots may distort cell-cell similarities in single-cell gene expression (36,37), we verified our observations on glycogene expression directly, without relying on UMAP latent embeddings.

### Glycopathway expression vary with tissue and cell type of origin

A rich diversity of glycan structures arise from sets of reactions operating together as “glycopathways”. To gauge the expression of these glycopathways in TS cells, we calculated the expression of glycopathways in the TS cells by taking the average expression of the genes from each glycopathway as delineated in the GlycoEnzOnto (see **Suppl. Table S3**). To further discern the patterns of glycopathway expressions, differential expression (DE) analysis was performed (**Fig. 2**). As illustrated in **Fig. 2A**, the DE analysis combined two information: zero-inflated mean expression of glycopathways for cells in each tissue type; and log-fold change (logFC) of the glycopathway mean expression value in a given tissue against cells in all other tissues. The intensity of green color in the DE heatmap plot informs the glycopathway expression while the border thickness indicates the logFC. In the example of enzymes involved in extension of the glycolipid (GSL) core and its diversification into ganglio-, lacto-, neolacto- and globo-series (**Fig. 2A**), we found a higher expression of relevant genes in eyes, trachea and small intestines compared to other tissues.

**Fig. 2B** visualizes the mean and differential expression of various glycopathways across epithelial cells found in multiple tissue-types. Here, glycopathways are grouped into three major functional categories: Core, Extension, and Terminal pathways (5). The results for three additional glycopathway groups: Core subclass, Nucleotide Sugar (NS) metabolism, and Degradation processes are provided in **Suppl. Fig S3A**. The same analyses are also performed for Endothelial, Stromal and Immune cell types, and the results are presented in **Suppl. Fig. S4A-S6A**. **Suppl. Table 4** provides the mean expressions of the glycopathways for the cell types in the TS dataset. The color scale bar for mean expression used in Fig. 2 and the aforementioned supplemental figures is the same, allowing direct comparison between the cell/tissue types. Additionally, the observations are independent of cell sample size since the mean expression of glycogenes in a system did not depend on either the number of cells in the tissues (𝜌 = −0.22, *p*-value = 0.30) or the number of cells of a given population (𝜌 = −0.15, *p*-value = 0.81).

The data present several striking observations. Notably, endothelial and stromal cells consistently manifested higher levels of glycopathway expression in comparison to epithelial and immune cells. Surveying the groups of glycopathways, the Core, NS metabolism, and Degradation groups had the highest mean glycogene expressions. These observations are in agreement with the expression analysis of glycogenes from these pathways in **Fig. 1E** and **1F**. The trend also underscores the broad functions that these groups of glycopathways have in terms of controlling global glycan turnover rates and pathway initiation steps. In general, the most highly expressed core gene set across all cell/tissue type belonged to the O-GlcNAc forming enzymes OGT and OGA. This highlights the importance of O-GlcNAc post-translational modification in regulating a broad swath of cellular signaling, transcription and disease processes (38). Besides these enzymes, we also observed consistent high expressions of several other core-pathways in diverse cell types, particularly those initiating GSL biosynthesis (*i.e.*, ‘GlcCer-series’) and those initiating the synthesis of N-linked glycans (*i.e.*, ‘Dolichol pathways’). In addition to the core Dolichol pathway, high gene expression was also observed for N-glycosylation processing enzymes that trim the initial dolichol precursor to enable protein folding and the biosynthesis of complex type structures. Finally, the prevalence of core-3 and core-4 O-GalNAc biosynthetic genes were largely restricted to the intestines, and this too is consistent with literature knowledge (39,40).

Glycosaminoglycans were expressed in stromal cells, particularly with respect to hyaluronan forming enzymes (HAS1, HAS2, HAS3) which were consistently high in fibroblasts and connective tissue in multiple organs. Among the rarer O-linked glycan modifications, enzyme contributing to O-mannosylation of cadherin superfamily (41), particularly TMTC1 was highly expressed in vascular endothelial cells (**Fig. S4A**). Not much is known about these pathways, but the measured high gene expression warrants additional investigation regarding its biological function. Among the enzymes involved in nucleotide biosynthesis, we noted high levels of UGDH (UDPGlcA_synth) and UXS1 (UDPXyl_synth) which are involved in the biosynthesis of starting materials that contribute to glycosaminoglycan biosynthesis. Enzymes in the biosynthesis of other nucleotide-sugar donors were also present in all cells, albeit at lower levels. Finally, several enzymes involved in lysosomal targeting and trafficking (lyso_target) of glycosidases were also highly expressed (**Suppl. Table. S4**).

Among the enzymes mediating glycan extension, high expressions were noted in pathways involved in the biosynthesis of Type-II lactosamine (Galβ1-4GlcNAcβ) and Type-III lactosamine (Galβ1-3GalNAcβ) chains across all cell and tissue-types, compared to Type-I lactosamine (Galβ1-3GlcNAcβ) chains. This is generally consistent with current biological knowledge related to the high abundance of Type-II lactosamine chains on N-/O-linked glycans and GSLs. Interestingly, the abundance of I-branching enzymes GCNT2 and 3 were restricted to specific tissue types supporting the emerging notion that such GlcNAcβ1-6 branching may have important biological functions for example in regulating cell growth and survival (42). Additional extension glycopathways that were highly expressed were involved in the N-glycosylation processing enzymes that are expressed in the endoplasmic reticulum (Ngly_trim) and the keratan sulfate extension enzymes (KS_poly). The epithelial cells of the liver, which is a major source of heparan sulfate biosynthesis, also had expression of HS extension genes at high levels.

The expression levels of chain terminating enzymes were often low and heterogenous, across cell-types. The only exception to this were the enzymes involved in protein tyrosine sulfonation (TyrP-synth) which were uniformly expressed at high levels in all cells and tissues. This is consistent with the ubiquitous nature this modification. Genes related to ABO antigen had the highest expression in epithelial cells (see **Suppl. Table. S4**) as these cells are a major source of blood group antigens. Finally, among the lowly expressed glycopathways, SDA antigen (Sda) biosynthetic enzymes were restricted to epithelial and immune cells while the enzymes forming sialyl Lewis-X epitope were dominant in T-cells, monocytes and neutrophil populations based on expression of the enzyme FUT7.

The logFCs for glycopathway expression are generally moderate, ranging from –0.81 to 1.37, suggesting the prevalence of similar pathways in different cell types. Among the pathways, Core and Extension groups display greater logFC magnitudes than those in the Terminal group suggesting heightened tissue-to-tissue variability. Similar to the Terminal group, the Core subclass, together with the NS and Degradation groups, exhibits only modest logFCs across different tissues. Upon examining individual tissues, cells in the eye, heart, and lung typically demonstrated higher overall glycopathway expressions when compared to other cells. In contrast, cells in the kidney and liver display relatively lower overall expressions. Interestingly, cells originating from tissues that are related to each other, such as large and small intestines, present similar expression patterns as shown by hierarchical clustering in **Fig. 2** and **Suppl. Fig. S3**. Overall, several of our computational predictions are consistent with literature and our analysis also suggest additional hypothesis that require experimental validation. To facilitate such comprehensive exploration of glycopathways, we have developed a webtool that is accessible via: http://vgdev.cedar.buffalo.edu/glycocarta/.

While DE analysis focuses on differences in mean gene expression, we complemented this with enrichment/depletion (ED) analysis. This analysis determines if a specific tissue are disproportionately enriched or depleted of cells that are expressing a given glycopathway— termed as “expressing cells”—when compared to the overall proportion of expressing cells in the entire TS dataset. Given that a number of glycopathways demonstrated low single-cell expressions, the fraction of expressing cells has been proposed as a better metric for gauging expression within a cell population (35). To this end, for every combination of glycopathway and tissue, we constructed a contingency table showing the count distribution of expressing/non-expressing cells for the glycopathway and of cells from/outside of the tissue (**Fig. 3A**, see Methods). Utilizing this table, we evaluated the log2 odds ratio (logOR). A logOR greater or lesser than 0 suggests that the particular tissue contains more or fewer expressing cells than expected based on the overall proportion of expressing cells in the TS dataset. In effect, a positive (negative) logOR indicates enrichment (depletion) of cells expressing the designated glycopathway in that tissue.

**Figure 3.**
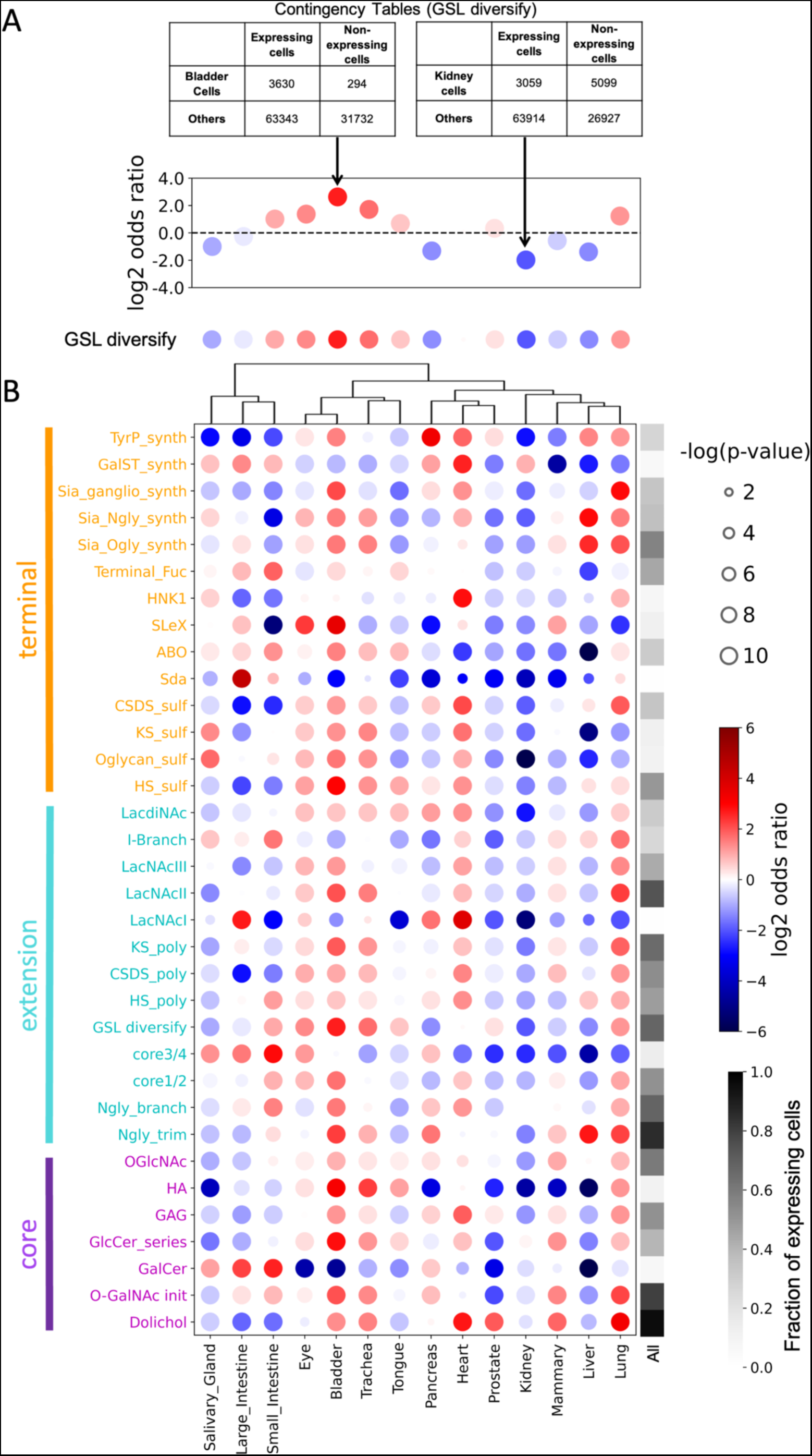
Enrichment-Depletion (ED) analysis of glycopathway in epithelial cells. (A) GSL (glycosphingolipids) diversify pathway is used to illustrate calculation scheme. ED analysis was performed by constructing the contingency table. The size of the dot represents the p-value of the Fisher exact test for significance, while the face color gives the sign of the log-odds ratio (logOR, blue: negative logFC and red: positive logFC). A negative logOR represents a depletion, while a positive logOR represents an enrichment of glycopathway in a tissue with respect to all other tissue with the same pathway. (B) ED of glycopathways in Core, Extension and Terminal groups for epithelial cells. The last column presents the fraction of expressing cells for each glycopathway among all epithelial cells using greyscale heatmap. This allows evaluation of how prevalent a given pathway is in epithelial cells compared to other pathways.

The results of the ED analysis as shown in **Fig. 3B** resonated well with the DE analysis findings. With the exception of the GalCer pathway in the kidney, pathways with positive (negative) logOR generally had positive (negative) log2FC. Also, there was a positive correlation between the logORs and the logFC (Pearson correlation 𝜌 = 0.61). In summary, the results of both DE and ED analysis highlight a moderate variability in glycopathway expressions across tissues, especially in the Core and Extension groups. Conversely, the Terminal, Core subclass, Nucleotide sugar and Degradation manifest higher variation in single-cell expression across different tissues. Additional wet-lab studies are warranted to determine how the variation in gene expression across tissue relate to tissue-specific glycan structure patterns.

### Transcriptional factors regulating glycosylation extracted from single-cell RNA-seq

Literature reported experimentally validated transcription factors (TFs) regulating glycogenes are few (5). To address this gap, we leveraged single cell transcriptomics data in the TS and the TF-gene binding interaction data in TFLink database (23). Our strategy involved evaluating mutual information (MI) of single-cell gene expression between every possible pair of transcription factor (TF) and glycogene. Here, mutual information (MI) gives a measure of how much the uncertainty in the expression of a glycogene is reduced given the corresponding expression data for a TF. Applying this strategy, the MI yielded an accurate prediction for TF-glycogene interactions, achieving an Area under Precision-Recall Curve (AUPRC) of 0.447, when compared to TF binding interactions sourced from the TFLinks (23). This accuracy outperformed both the pairwise Pearson correlation (AUPRC = 0.367) and the Spearman rank correlation (AUPRC = 0.396). All of the above interaction scores: MI, Pearson correlation, and Spearman rank correlation, surpass the performance of a random predictor (AUPRC = 0.318). This outcome suggests that single-cell gene expression data may be used to infer TF-glycogene interactions.

Our single-cell TF-glycogene evaluation facilitates the identification of regulatory modules (RMs) of glycosylation. In this context, a RM refers to a set of glycogenes whose transcription is controlled by a shared regulatory program (43). To this end, we performed a hierarchical clustering of glycogenes using their log1p of MI scores with 1184 TFs (i.e., log(1+MI)). The clustering reveals five RMs, as depicted in **Fig. 4A** (see **Suppl. Table S5** for glycogene membership in clusters). Subsequent analysis using Fisher’s exact test linked each cluster with specific glycopathway classes (two-sided Fisher’s exact test, Benjamini-Hochberg adjusted p-value < 0.1; see Methods). The odds ratios presented in **Fig. 4B** (see also **Suppl. Fig. S7**) indicate that different RMs are associated with distinct classes of glycopathways, implying a shared transcriptional regulatory program among glycogenes from the same pathway class. We also curated a ranked list of TFs for each RM (see **Suppl. Table S6**), providing insights into potential regulatory factors. To explore the TF-glycogene analysis more fully, we developed a web-tool: http://vgdev.cedar.buffalo.edu/glycotf/.

**Figure 4.**
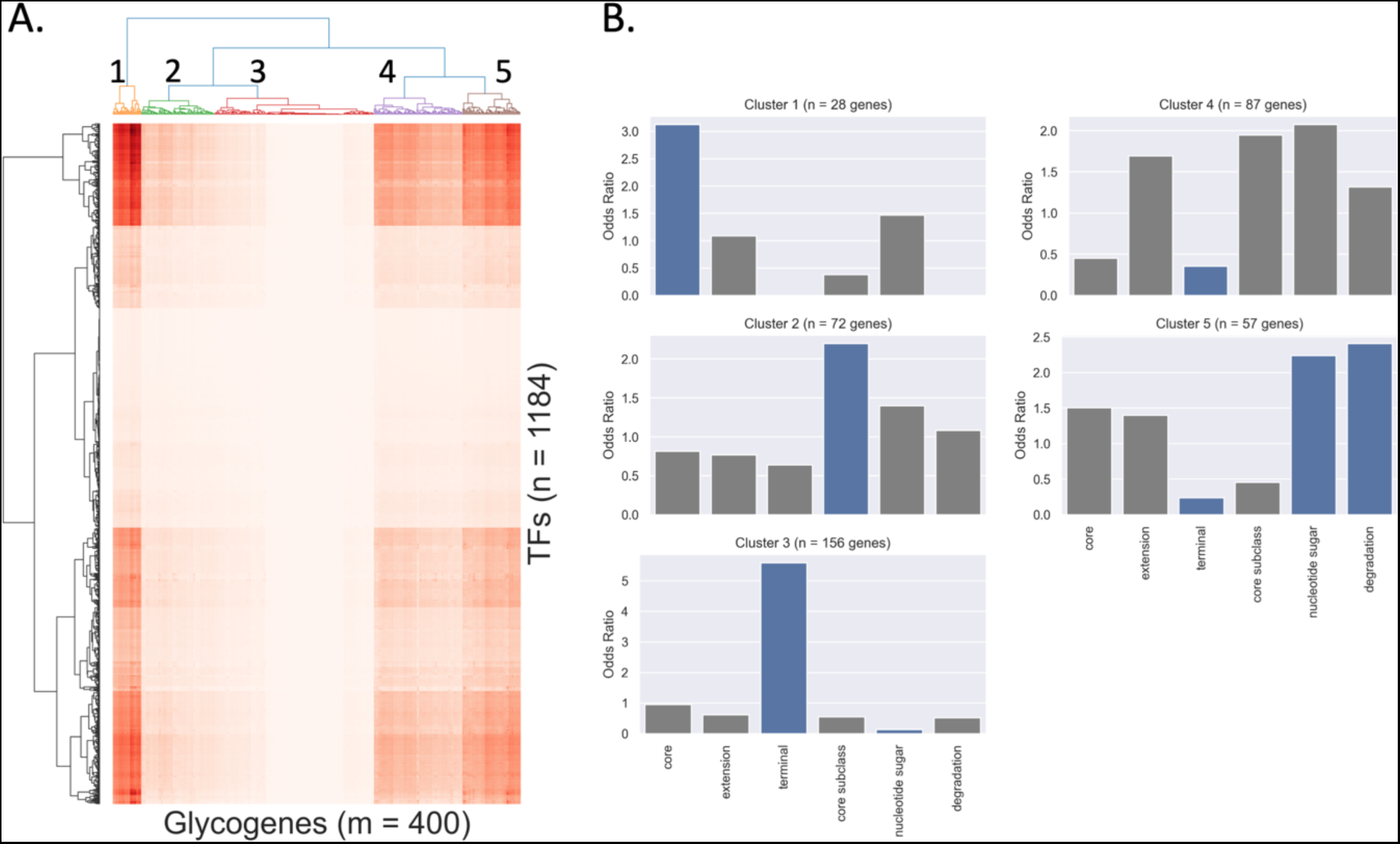
Glycosylation Transcription Factor Analysis. (A) Hiearchical clustering of glycogenes based on the log1p of MI scores of TF-glycogene (*i.e.*, (log(1+MI)). The numbers indicate cluster labels. (B) Enrichment analysis of glycogenes in each cluster for glycopathway classes. Blue bars indicate those with Benjamini-Hochberg adjusted *p*-value < 0.1 (two-sided Fisher exact test).

The first RM (Cluster 1) is strongly associated with the synthesis of Core glycan structures. The glycogenes in this cluster encode enzymes involved in the formation of the oligosaccharyltransferase (OST) complex (i.e., dolichol pathway): DAD1, DDOST, RPN1, RPN2, and STT3B; enzymes involved in initial processing of N-glycans: PRKCSH and GANAB; glycoprotein folding chaperones: CANX, CALR, ERLEC1, HSP90B1, HSPA5, IGF2R, LMAN2, OS9, SE1L; O-GlcNAc biosynthesis enzymes: OGT and OGA; and other high abundance genes: B4GALT1, GPI, M6PR, and MGAT1. The second RM (Cluster 2) is connected to a multitude of processes involves in the Core subclass (see also **Supp. Fig. S7**). Specifically, glycogenes in Cluster 2 are involved in a number of glycopathways responsible for the biosynthesis of GAG and lipid-linked oligosaccharides, and the O-linked glycan post-translation modification such as POFUT and POMT genes. A number of transporter genes also belong to Cluster 2.

The third RM (Cluster 3) is significantly enriched for glycogenes involved in the Terminal class. More specifically, cluster 3 includes UDP-Glucuronosyltransferase family genes that are involved in the glucuronidation that enable drug metabolism and also metabolism of pollutants, bilirubin, androgens, estrogens, mineralocorticoids, glucocorticoids, fatty acid derivatives, retinoids, and bile acids. This cluster also comprises a number of sulfotransferases that modify both GAGs and glycoproteins; a majority of genes (8 out of 11) that participate in the terminal protein fucosylation; and several members of the sialyltransferase family (10 out of 20). Importantly, the Terminal class is under-represented in all clusters besides Cluster 3 (odds ratio < 1), suggesting that these processes are under a distinct transcriptional regulatory program than the others.

The fourth RM (Cluster 4) is intermixed with glycogenes from the NS metabolism, Core Subclass, and Extension groups, but none of these classes crossed the statistical significance cutoff (Benjamini-Hochberg adjusted p-value < 0.1). Important genes involved in the synthesis of nucleotide sugars, such as GALT, GALK2, GALE, PGM1-3, and GMPPA/B, belong to this RM. Another prominent feature of Cluster 4 is the presence of genes involved in the initiation of heparan sulfate and chondroitin sulfate biosynthesis including B4GALT7, FAM20B, B3GALT6, CHPF, CHPF2, EXT1, and EXT2. This cluster also includes genes involved in GPI anchor biosynthesis (PIGC, PIGG, PIGH, PIGK, PIGN, PIGS, PIGX) and in the modification of N-glycosylation, specifically in terminal sialylation and core-fucosylation (ST6Gal1 and FUT8). The reason why a single cluster of TFs would regulate a diverse group of pathways remains to be studied in literature.

The last RM (Cluster 5) is strongly linked to NS metabolism and Degradation classes. Genes involved in NS biosynthesis in this cluster comprise CMAS, DPM1, DPM2, GALK1, GFPT1, GFUS, GNPDA1, PAPSS, UAP1 and UGDH. The cluster also includes a set of genes involved in the degradation processes, such as CTBS, CEMIP2, GLB1, GNS, GUSB, HEXA, HEXB, HGSNAT, IDS, NEU1, and FUCA2. Except for collagen degradation, all glycopathways in the Degradation class are over-represented in this RM (odds ratio > 1, see **Suppl. Fig.** S7).

## DISCUSSION

A key contribution from this study is the description of the broad landscape of glycoEnzymes and glycopathways in normal human cell and tissue types. The findings are in broad agreement with the recent work by Joshi and colleagues (34) that showed that core enzymes are more ubiquitously expressed among cells than enzymes that modify terminal glycan residues which cater to more specialized functions. However, it is important to note differences in the study design as the glycogene set used in this work is considerably larger (224 in Joshi vs. 400 in this work), owing to the inclusion of glycosidases, transporters and other regulators of carbohydrate biosynthesis. In addition, the focus on glycopathways and well-defined ontologies represents a step away from the previous approach. Importantly, we also analyze single-cell gene expression data directly without pseudobulking. The use of differential expression (DE) and enrichment-depletion (ED) analysis, as opposed to using inter-quartile distances to judge the importance of specific glycogenes, is another difference. Finally, our study presents the first detailed analysis of TF-glycogene relations using single-cell gene expression data. This reveals the possibility that distinct TF regulatory modules control various aspects of mammalian glycosylation.

While our study presents a broad analysis of human glycosylation pathways, it is not without limitations. At the sequencing depth employed in the TS, out of the anticipated ∼220 glycogenes expressed human cells, less than one fifth are detected in a typical cell in this dataset. While this low detection is comparable with other PC genes, such high data sparsity impedes data analysis such as single-cell clustering. The underlying TS data also does not include key organs like the brain which have distinct glycosylation profiles compared to other organs. Further data analysis is thus required to integrate the findings of this work with brain initiatives and related activities (44). In addition, while our analysis captures the nuances of gene expression patterns related to glycosylation, additional wet-lab studies are needed to extrapolate these findings to glycan structures on the individual cell types and related functional outcomes. Such an endeavor requires the development of novel technologies to measure glycoenzyme activity in greater depth at single cell level and parallel development of glycomics analysis. Such effort would provide understanding of the myriad of steps regulating glycosylation (transcription, translation, biosynthetic reactions) and their impact on cell function. Thus, at this time, the observed variations in gene expression described in this manuscript only describe a portion of the factors affecting cellular variations in the glycome. Direct validation, such as through glycomics analysis at single-cell level and CRISPR technology based molecular screens are paramount in solidifying our inferences and bridging the gap between gene expression and functional glycan structures on proteins.

## DATA AVAILABILITY

The data underlying this article are available in Figshare, at https://doi.org/10.6084/m9.figshare.14267219. The source codes are available in Github, at http://www.github.com/cabsel/glycots.

## SUPPLEMENTARY DATA

Supplementary Data are available online.

## ACKNOWLEDGMENTS

This work was supported by the National Institutes of Health (grant #3R01HL103411-10S1).

S.V. was partially supported by SUNY Multidisciplinary Small Team Award (grant #201047.2). Funding for open access charge: National Institutes of Health.

