## Supplementary material for "Cell and tissue-specific glycosylation pathways informed by single-cell transcriptomics": Suppl. Fig.

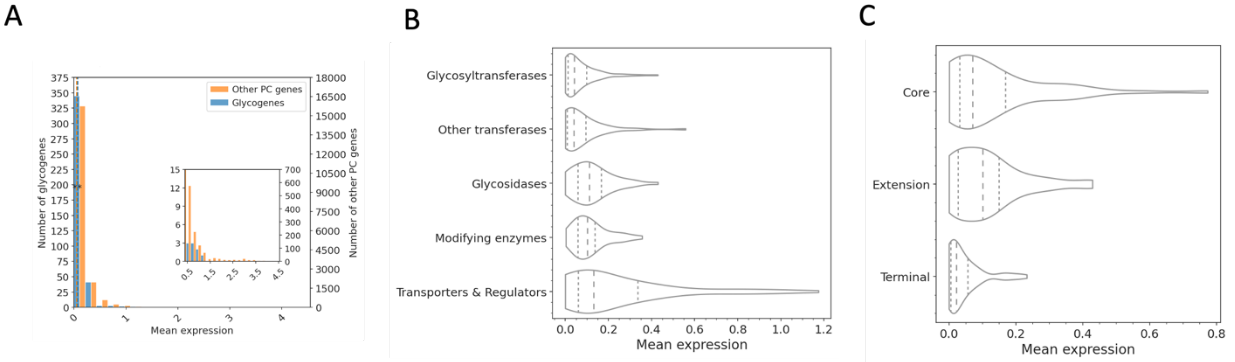


**Fig. S1.** (A) Distribution of protein coding (PC) genes and glycogenes based on their mean expression in the single cells of Tabula Sapiens dataset. The median value for glycogenes is higher than that for other PC genes (p-value = $1.36 \times 10^{-3}$, two-sided Wilcoxon ranksum test). (B) Distribution of mean expression of glycogenes. Glycogenes are grouped based on their biological functions as defined in GlycoEnzOnto: Glycosyltransferases, Other transferases, Glycosidases, Modifying enzymes, and Transporters and Regulators. (F) Distribution of single cell expression of glycosyltransferases in Core, Extension and Terminal groups (see Supplementary Table S3).

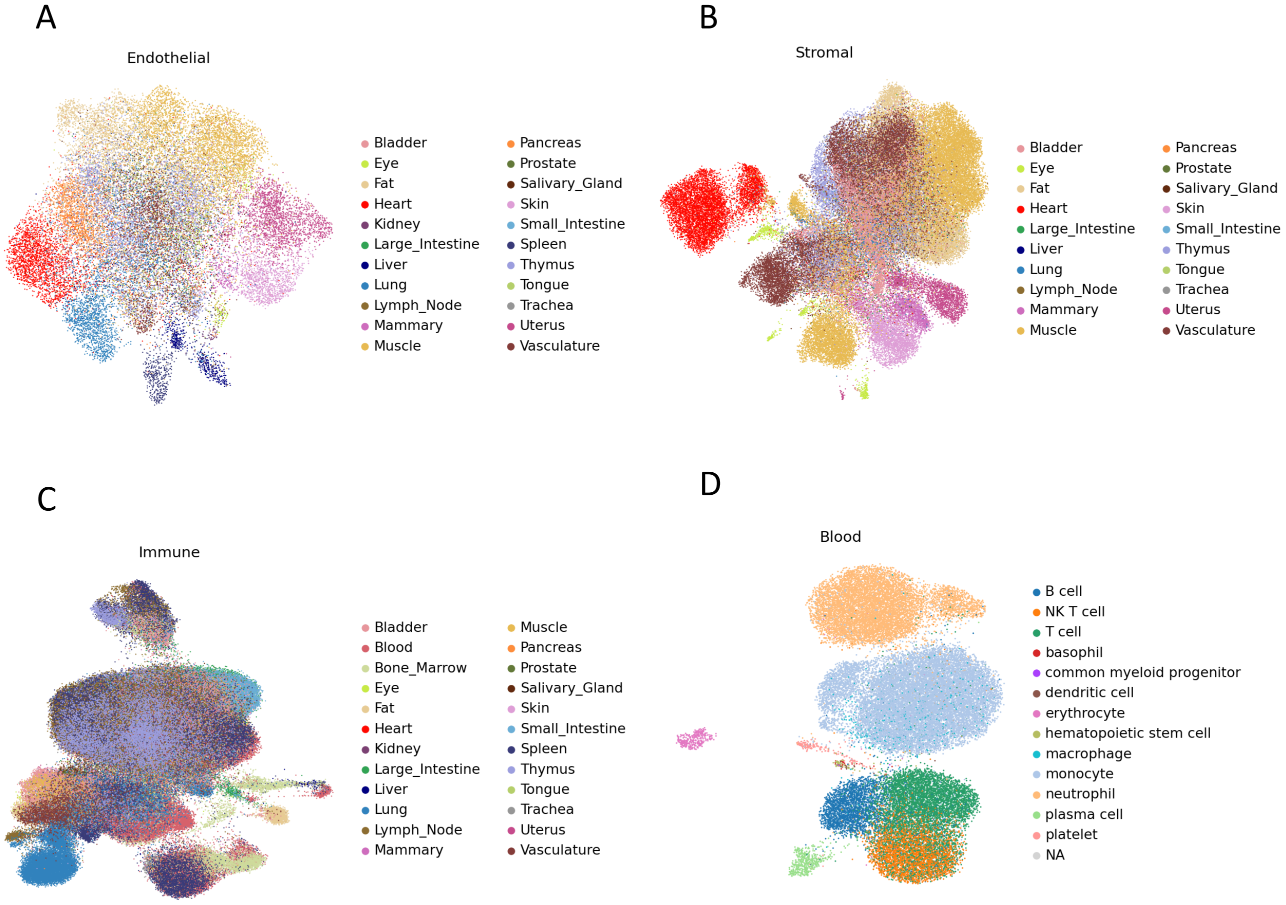


**Fig. S2.** UMAP visualization of scVI (single-cell Variational Inference) latent embedding of glycogene expression in (A) endothelial, (B) stromal, and (C) immune cells and (D) blood cells (including erythrocytes and platelets).

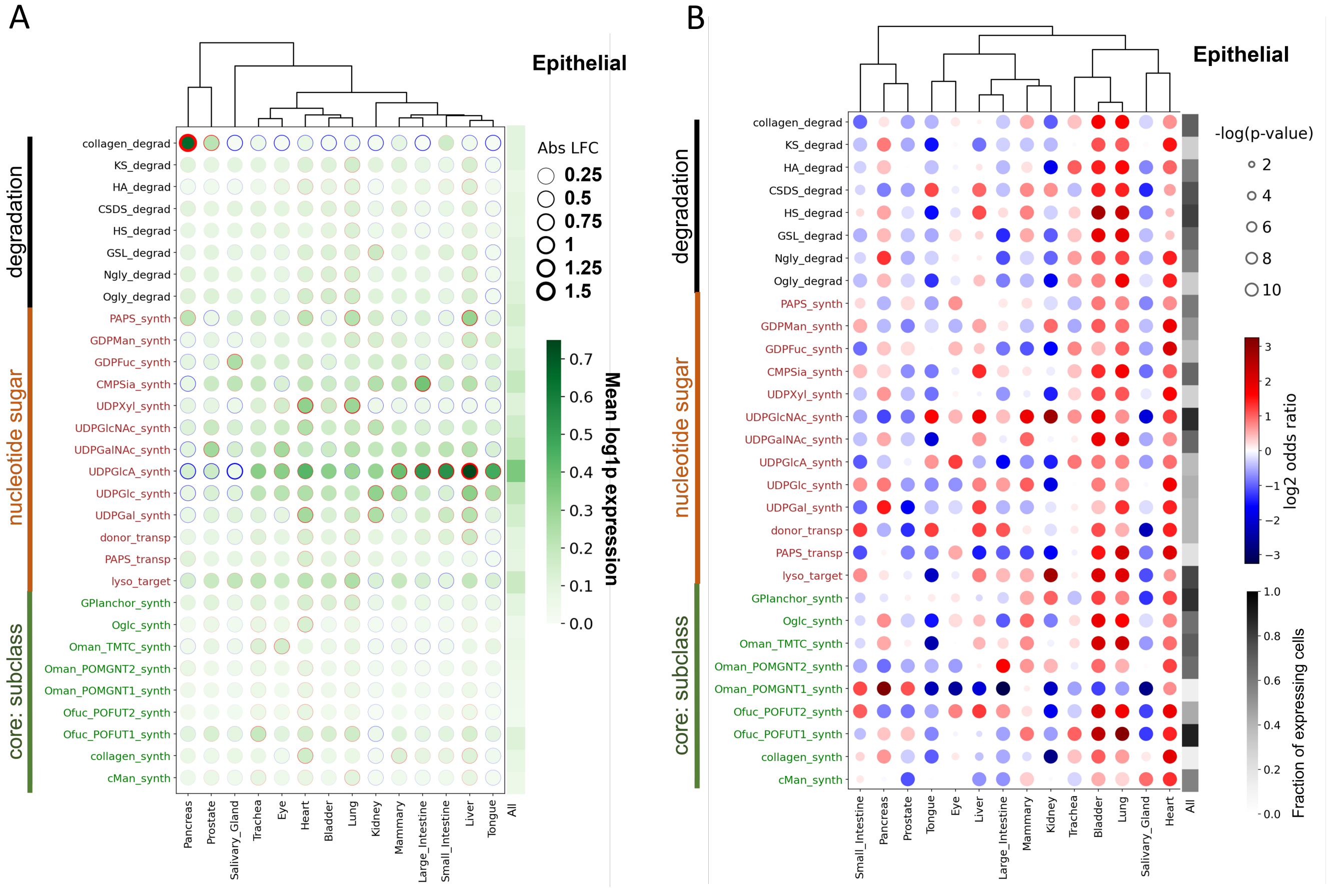


**Fig. S3.** (A) Differential expression (DE) analysis of glycopathway in epithelial cells. DE of glycopathways is presented for selected Core subclass, Nucleotide sugar and Degradation pathways for epithelial cells. The greenscale heatmap in the last column presents the mean glycopathway expression among all epithelial cells. (B) Enrichment-Depletion (ED) analysis of glycopathways in Core subclass, Nucleotide sugar and Degradation groups for epithelial cells. The size of the dot represents the p-value of the Fisher exact test for significance, while the face color gives the sign of the log-odds ratio (blue: negative logFC and red: positive logFC). A negative logOR represents a depletion, while a positive logOR represents an enrichment of glycopathway in a given tissue with respect to all other tissues with the same pathway. The last column presents the fraction of expressing cells for each glycopathway among all epithelial cells using greyscale heatmap.

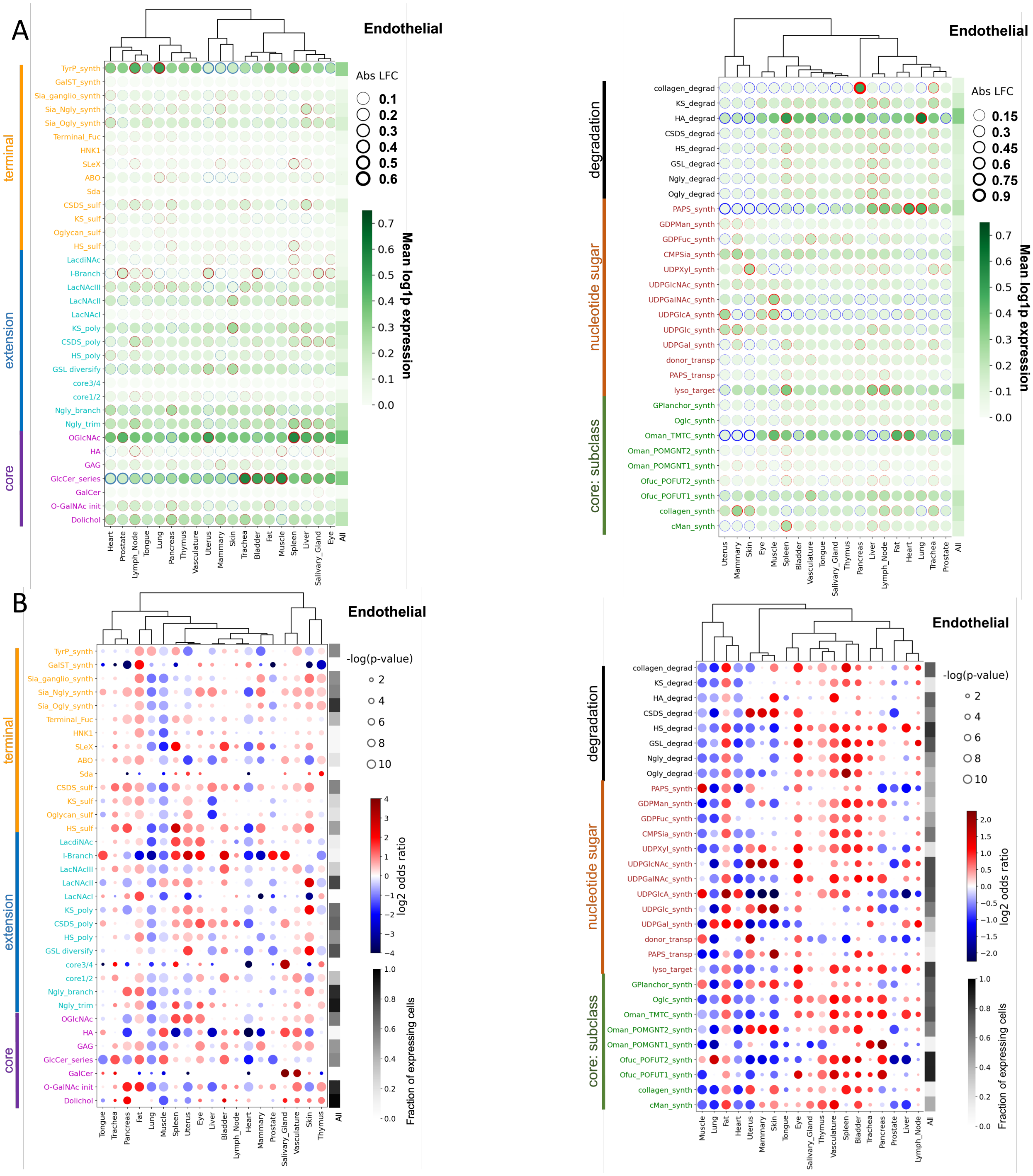


**Fig. S4.** (A) Differential expression (DE) analysis of glycopathway in endothelial cells. DE of glycopathways is presented for Core, Extension, Terminal, Core subclass, Nucleotide sugar and Degradation pathways for endothelial cells. The greyscale heatmap in the last column presents the mean glycopathway expression among all endothelial cells. (B) Enrichment-Depletion (ED) analysis of glycopathways in Core, Extension, Terminal Core subclass, Nucleotide sugar and Degradation groups for endothelial cells. The size of the dot represents the p-value of the Fisher exact test for significance, while the face color gives the sign of the log-odds ratio (blue: negative logFC and red: positive logFC). A negative logOR represents a depletion, while a positive logOR represents an enrichment of glycopathway in a given tissue with respect to all other tissues with the same pathway. The last column presents the fraction of expressing cells for each glycopathway among all endothelial cells using greyscale heatmap.

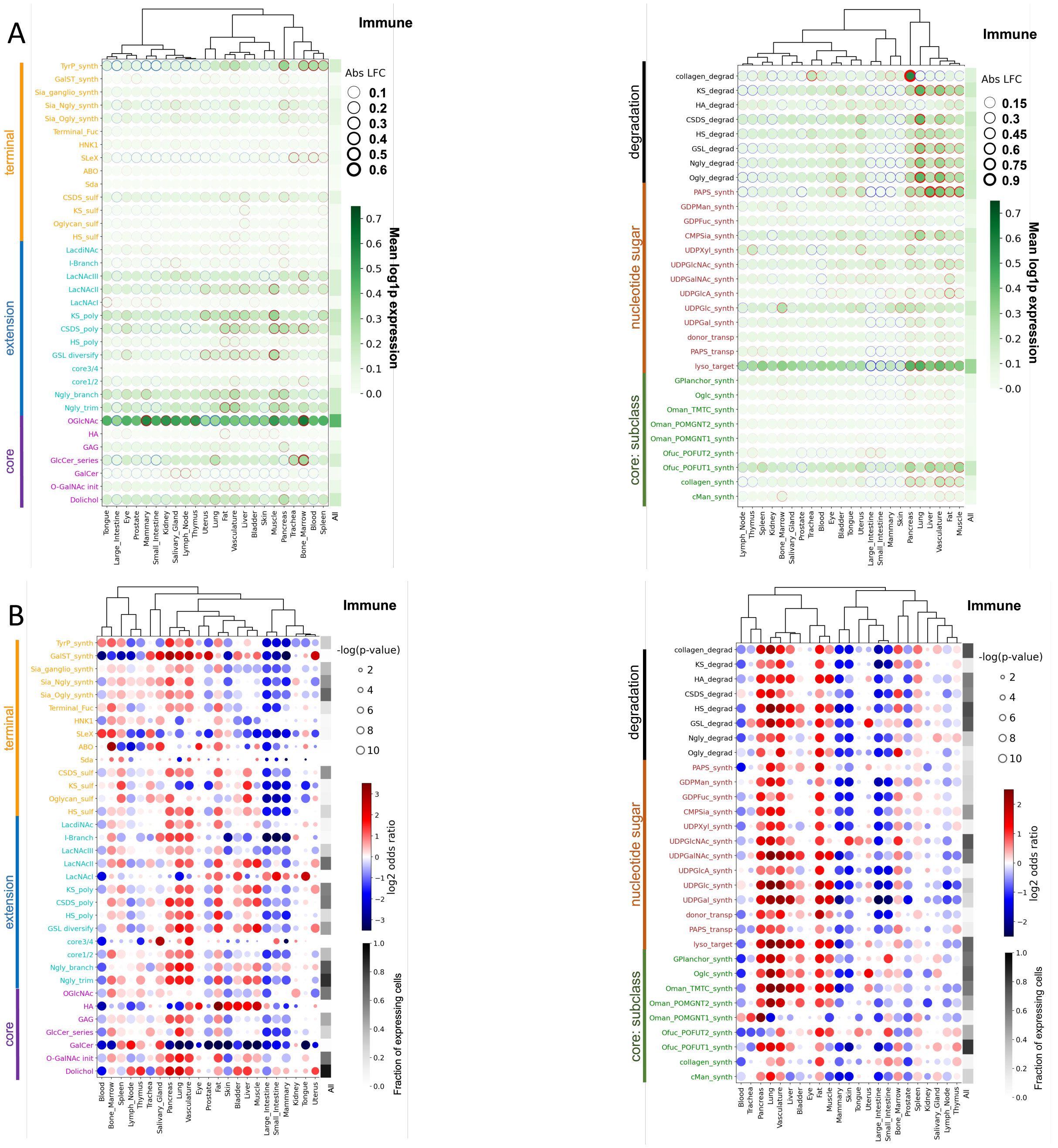


**Fig. S5.** (A) Differential expression (DE) analysis of glycopathway in immune cells. DE of glycopathways is presented for Core, Extension, Terminal, Core subclass, Nucleotide sugar and Degradation pathways for immune cells. The greyscale heatmap in the last column presents the mean glycopathway expression among all immune cells. (B) Enrichment-Depletion (ED) analysis of glycopathways in Core, Extension, Terminal Core subclass, Nucleotide sugar and Degradation groups for immune cells. The size of the dot represents the p-value of the Fisher exact test for significance, while the face color gives the sign of the log-odds ratio (blue: negative logFC and red: positive logFC). A negative logOR represents a depletion, while a positive logOR represents an enrichment of glycopathway in a given tissue with respect to all other tissues with the same pathway. The last column presents the fraction of expressing cells for each glycopathway among all immune cells using greyscale heatmap.

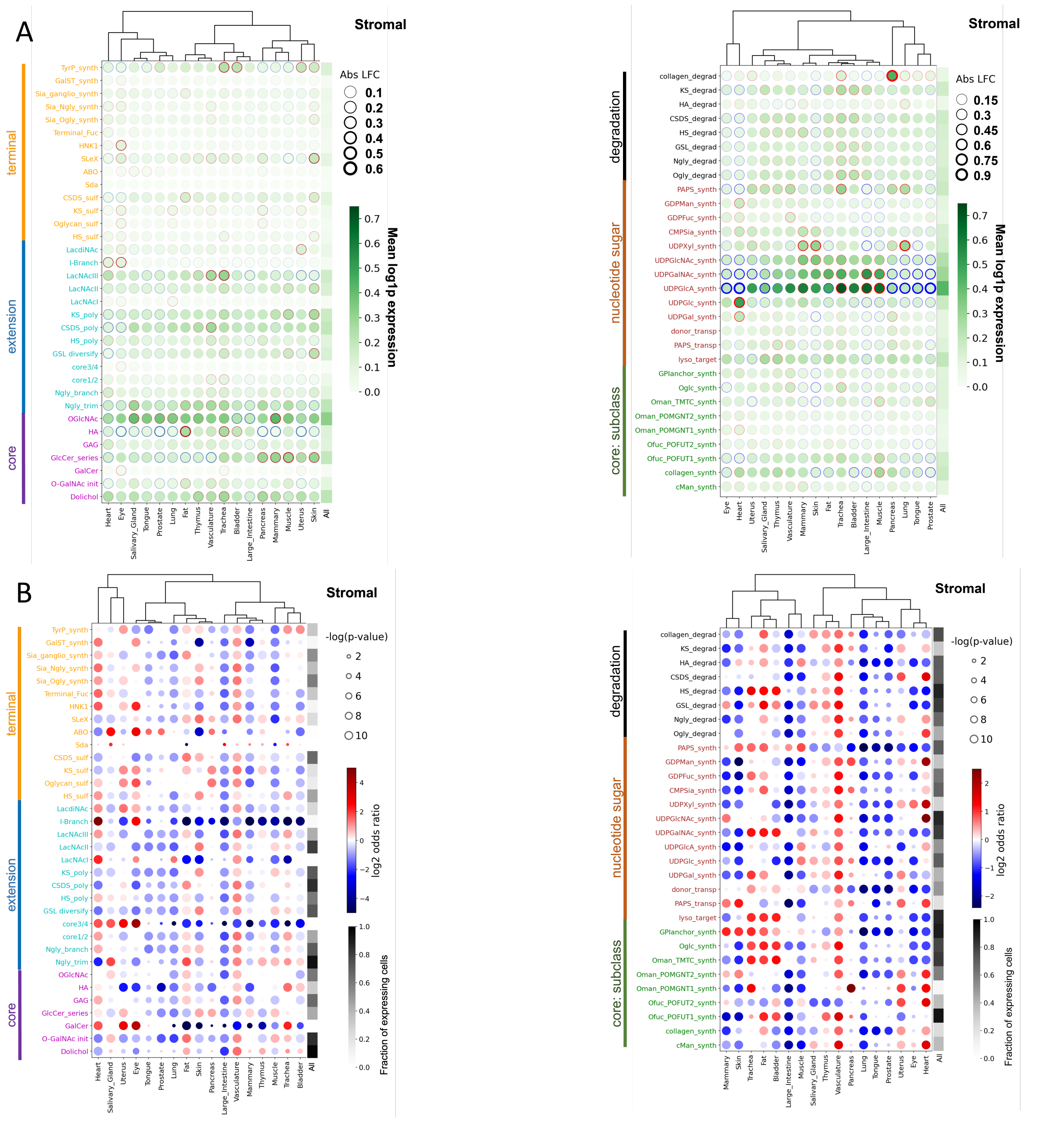


**Fig. S6.** (A) Differential expression (DE) analysis of glycopathway in stromal cells. DE of glycopathways is presented for Core, Extension, Terminal, Core subclass, Nucleotide sugar and Degradation pathways for stromal cells. The greyscale heatmap in the last column presents the mean glycopathway expression among all stromal cells. (B) Enrichment-Depletion (ED) analysis of glycopathways in Core, Extension, Terminal Core subclass, Nucleotide sugar and Degradation groups for stromal cells. The size of the dot represents the p-value of the Fisher exact test for significance, while the face color gives the sign of the log-odds ratio (blue: negative logFC and red: positive logFC). A negative logOR represents a depletion, while a positive logOR represents an enrichment of glycopathway in a given tissue with respect to all other tissues with the same pathway. The last column presents the fraction of expressing cells for each glycopathway among all stromal cells using greyscale heatmap.

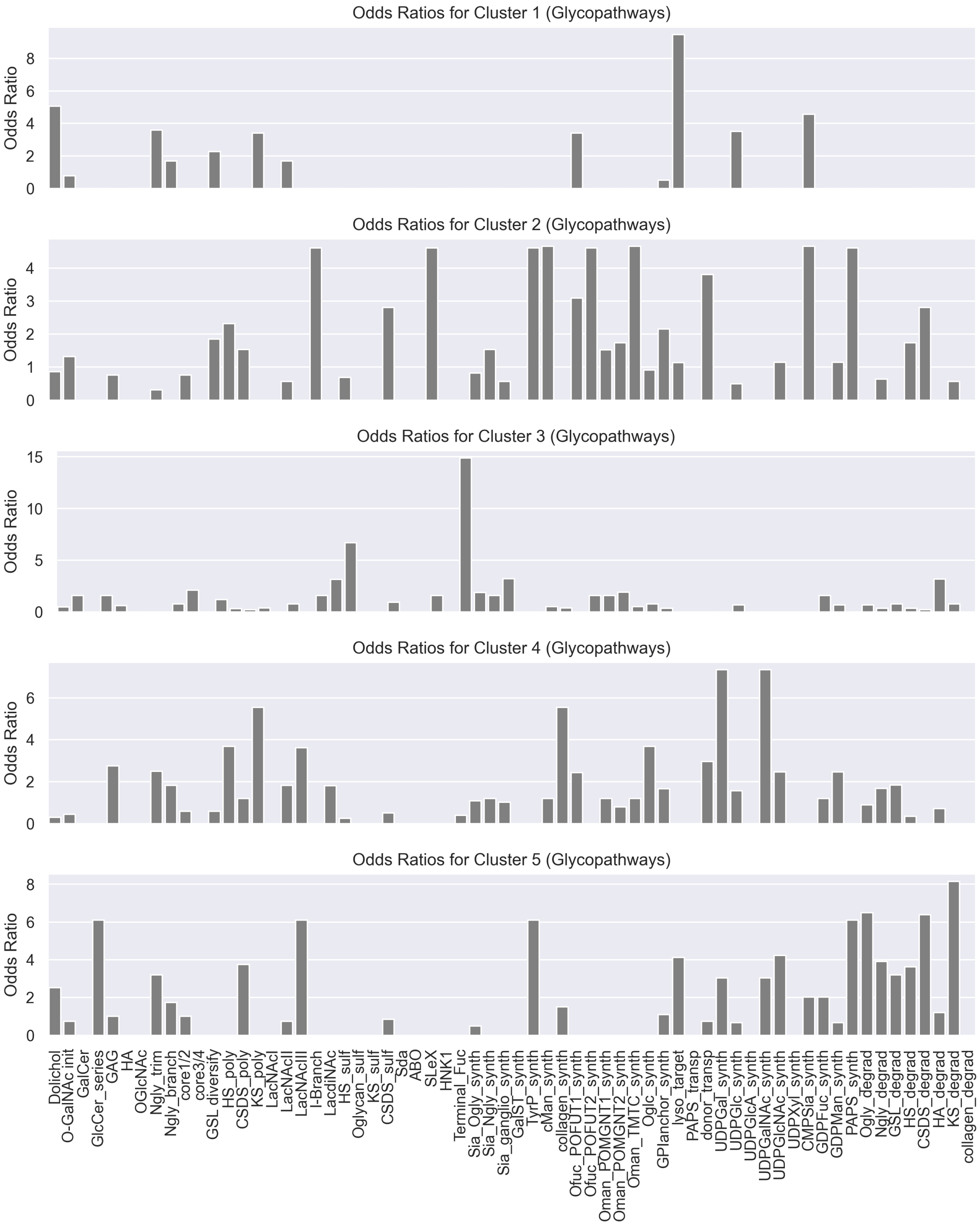


**Fig. S7.** Enrichment analysis of glycogenes in each cluster for different glycopathways. None of the glycopathways are significantly enriched (adjusted p-value > 0.1, two-sided Fisher exact test with Benjamini-Hochberg correction).
